# Temperature and management reduce the abundance of the invasive Argentine ant (*Linepithema humile*)

**DOI:** 10.64898/2026.10.08.756264

**Authors:** Daniel Sánchez-García, Xim Cerdá, Elena Angulo

## Abstract

**BACKGROUND:** The Argentine ant (*Linepithema humile*) is among the world’s worst invasive species. In the Doñana Biological Reserve (Spain), it is associated with human structures that provide the mild, moist conditions it needs within the hot, dry scrubland. We evaluated a habitat-modification treatment, predicting that removing this suitable habitat would eradicate the invader locally. Over an 18-year period, we monitored ant communities at a treated and an untreated site, at different distances from the human structures, before and after treatment, and analysed how temperature and worker numbers affected encounters at baits between the Argentine ant and native ants.

**RESULTS:** Distance to the human structures shaped ant communities at both sites, but only the treated site changed after treatment. There, the Argentine ant decreased close to the human structure and almost disappeared farther from it, whereas the endemic *Cataglyphis tartessica* and *C. floricola* recovered, especially close to the structure. At baits, the outcome of encounters depended mainly on how temperature changes favoured each species relative to its maximum activity temperature: *Cataglyphis* spp. displaced the Argentine ant in the morning, as temperature increased, and the Argentine ant recovered the baits in the late afternoon. Numerical advantage favoured only the Argentine ant.

**CONCLUSION:** Habitat modification reduced the Argentine ant and allowed endemic ants to recover, but did not eradicate the invader, which persisted at low abundance 13 years after treatment. In protected areas, habitat management should be combined with eradication, monitoring and prevention of spread.

## 1 Introduction

One of the major drivers of biodiversity loss is invasive alien species, which are introduced into a novel range where they establish, spread and cause impacts.^1,2^ Ecological impacts of invasive species range from the extinction of native species to the disruption of ecosystem structure and function.^3–6^ Moreover, beyond ecological impacts, invasive species also affect multiple sectors generating massive and growing economic costs.^7^ Thus, managing invasive species remains a major challenge for society.^8^

Management actions follow the stages of invasion, theoretically beginning with the prevention of introduction and other biosecurity actions, followed by rapid eradication at early-invasion stages, and ending with long-term management such as control and containment at late-invasion stages.^8,9^ However, in practice, management is often reactive, triggered only after the impacts are detected.^10,11^ Moreover, delays in perceiving impacts frequently postpone management responses - if they are implemented at all.^11,12^ This can occur locally due to different perceptions of the impacts, limited information, inadequate invasive species prioritisation, or a shortage of funding for implementing management actions.

Ants have been introduced worldwide and continue to spread globally at high rates,^13,14^ with over 200 species introduced outside their native range. Many of them are cosmopolitan species closely associated with human buildings or infrastructures.^15^ Several cause negative impacts on ecosystems by displacing native ants, disrupting ecosystem processes and homogenising community assemblages.^16–18^ Beyond the environmental damage, invasive ants also impose major costs on socio-economic sectors including agriculture, forestry, public health and social welfare.^19^

Although not the costliest, the Argentine ant, *Linepithema humile* (Mayr, 1868), is listed among the world’s worst invasive species by the International Union for Conservation of Nature due to its severe effects on biodiversity.^19–21^ As one of the best studied invasive ant species,^21,22^ we know that in climatically suitable areas the success of this ant requires, among other factors, replacing competing native ants^23,24^ and co-opting mutualisms with homopterans.^25^ Successful invasions can disrupt key ecosystem processes such as pollination,^26^ seed-dispersal,^27,28^ or predator-prey interactions.^29–31^

The successful establishment of this invader is influenced by a combination of the abiotic and biotic conditions of the recipient ecosystem and the traits of the invader.^1^ The Argentine ant is native to the Paraná river basin, spanning eastern Bolivia, southern Brazil, Paraguay, Uruguay and northern Argentina.^21^ Outside its native range, it primarily invades mainly Mediterranean-type climatic areas.^32^ Indeed, its expansion is limited by precipitation^33–36^ as well as temperature, given that this species cannot tolerate extreme heat (thermal limit CT_max_: 38–40°C).^37^

In Europe, the Argentine ant occupies mainly coastal areas where it benefits from the mild conditions matching its native climate.^32,38,39^ In the coastal area of the Doñana Biological Reserve (the core of Doñana National Park) the species was introduced in the 1970s with the material used to rebuild houses following the National Park’s declaration. Following a decade of population growth around these human structures, the Argentine ant was first detected in natural habitats in 1981. However, research only began in the late 1990s, and monitoring was initiated in the 2000s, given its expansion in the Reserve, where it preferentially occupies scattered cork oak trees in the scrubland matrix or pine forests.^40^ Nevertheless, the high temperature and low humidity of the scrubland matrix, which constitutes the main terrestrial habitat of the Reserve, currently limit its spread to other habitats.^41^

Here we assessed the effectiveness of a habitat-modification treatment to control or contain the Argentine ant in the Doñana Biological Reserve. We predicted that the removal of the available suitable habitat would be sufficient to eradicate the species locally. To test this, we monitored the ant community over an 18-year period across two invaded sites featuring human structures surrounded by the scrubland matrix. The treatment consisted of replacing a concrete fence with a wooden one at one site - thereby eliminating the moist, shaded microhabitat that supported ant nests - while the second site served as an untreated control. We evaluated the recovery of the native ant community by analysing co-occurrence patterns, community structure, and the abundance of native endemic species over time. Additionally, we examined the influence of temperature on bait replacement, dominance hierarchies and daily foraging activity.

## 2 Material and methods

### 2.1 Study site

The field study was conducted at the Doñana Biological Reserve (37°01’N, 6°33’W; Doñana National Park, SW Spain). Doñana is located at the mouths of the Guadalquivir and Guadiamar rivers and comprises three primary Mediterranean ecosystems: coastal dunes, riverine marshlands and the scrubland between them. Within the scrubland, savine juniper forest (*Juniperus phoenicea* Linnaeus), pine forest (*Pinus pinea* Linnaeus) and isolated cork oak trees (*Quercus suber* Linnaeus) are scattered within the scrubland matrix.

The Doñana Biological Reserve hosts over 30 ant species, including two Andalusian endemic *Cataglyphis* species (*Cataglyphis floricola* Tinaut, 1993 and *Cataglyphis tartessica* Amor & Ortega, 2014).^42^ The Argentine ant likely entered the Park from the adjacent coastal village of Matalascañas,^38^ initially invading park buildings.^43^ In the 1970s, it was introduced to the core of the Reserve during the restoration of El Palacio de Doñana (in the centre of the Reserve), and by the 1990s it had spread into natural habitats and to another building 4.5 km away, “El Martinazo”^23,43^ (Fig. 1a). Currently, the ant occupies both human structures and natural habitats, presenting a patchy distribution with a preference for pine forests and scattered cork oak trees, while avoiding the surrounding hot and dry scrubland.^40,41^

**Figure 1.**
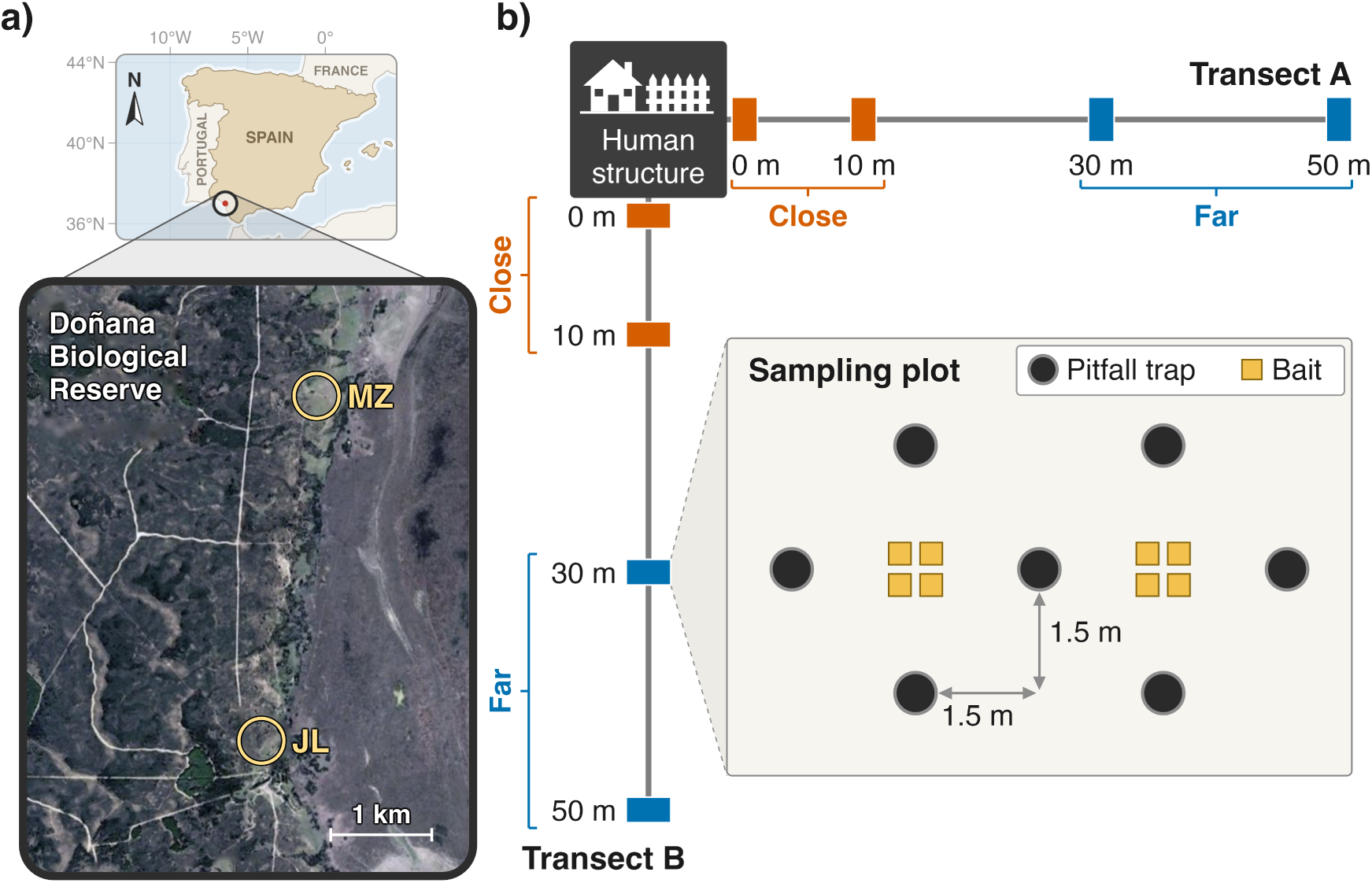
(a) Location of the two study sites in the Doñana Biological Reserve (SW Spain): the treated site (JL, “Jaulón del Lince”) and the untreated site (MZ, “El Martinazo”). (b) Sampling layout showing transects and plots at each site.

### 2.2 Management implemented

In 2000, two study sites were selected to investigate ant communities and their response to management actions (Fig. 1a). Both sites consist of human structures, where the Argentine ant was inadvertently introduced probably with food or material distribution from el Palacio. The first site is called “Jaulón del Lince” (JL, herein), and consists of a concrete fenced area isolated in the scrubland matrix, where during the 70’s and 80’s wild carnivores were maintained and fed. The Argentine ant was first detected in a cork oak tree near this area in 1981^23^ (Tinaut A, pers. comm.). The second site is the “Martinazo” (MZ, herein) house, an intermittently inhabited construction. It was first seen invaded by the Argentine ant in 1991, when it was inhabited.^43^ Both sites are surrounded by a matrix of scrubland, which is not suitable for the Argentine ant.^40,41^ In 2005, a management action was carried out in JL site in order to substitute concrete for wood in the fence, in order that a natural control of the Argentine ant could occur because of the lack of a suitable refuge against the hot and dry conditions of the scrubland.^40,41^ We compared these two sites over time, using the managed JL site as the treated and MZ as the untreated site.

### 2.3 Sampling ant communities over time before and after treatment

To evaluate ant presence, abundance, and activity at each site, we established two 50-m transects originating at the human structure (fence or house) and set at a 90° angle to each other (Fig. 1b). Along each transect, we set four sampling plots at distances of 0 m, 10 m, 30 m and 50 m from the human structure (Fig. 1b). Each plot consisted of seven pitfall traps in a double-diamond layout (1.5 m spacing) and, except the plots at 50 m, eight baits divided into two groups of four inside each diamond (Fig. 1b). Pitfall traps were plastic vials (6 cm diameter, 7 cm depth) filled with soapy water and left active for 24 h. All collected ant specimens from pitfall traps were preserved in 70 % ethanol and identified to the species level in the laboratory. Pitfall sampling was conducted from 2000 to 2005, 2008 to 2011, and in 2013 and 2018 (except at the treated site in 2000).

Ant foraging activity was measured at baits over the course of one day in each habitat using the groups of four baits of each study plot at 0, 10 and 30 m (Fig. 1b). Baits consisted of small plastic cards with four different food rewards (cheese, chorizo, biscuits, and water-diluted honey). The distance between the cards within each group was 25 cm. Baits were placed at 07:00h, and the first bait observation took place at 08:00h. Each hour from sunrise to sunset (08:00 until 23:00), the number of workers of each ant species feeding at each bait was recorded. Ground surface temperatures near the baits were measured with HOBO data loggers. For each group of bait records, we assigned the temperature value recorded by the sensor. Distances between baits and temperature sensors were 1.5 m. Ant activity was surveyed at both sites in July 2001 and July 2003. The untreated site was also surveyed in July 2000, before the management was carried out. The plots at 50 m had no baits, so the bait records of the far category correspond only to the plots at 30 m.

### 2.4 Temperature differences between sites

Differences in bait temperature between the two distance categories (close vs far) and between sites (treated vs untreated) were tested using Welch’s two-sample t-tests using the function *t.test()*, and, as a non-parametric alternative, with Wilcoxon rank-sum tests using the function *wilcox.test()*, with the normal approximation with continuity correction because of tied values. Both functions belong to the R package *stats*, included in R v4.5.0.^44^ Only temperature data from bait stations recorded during 2001 and 2003 were included in these analyses. Because a single data logger recorded the temperature of all the baits of a plot, the same reading was assigned to all the ant records of that plot in a given hour. Therefore, to avoid pseudoreplication, the tests were based on one temperature reading per plot and hour (n = 351 readings).

### 2.5 The success of the Argentine ant at baits

Bait records were first used to estimate the maximum activity temperature (MAT) of each species, defined as the temperature at which a species forages most actively at baits.^45^ For each species, temperatures recorded at baits occupied by at least one worker were grouped into 1 °C classes, and MAT was calculated as the mean temperature of the records in the class with the most records (modal class). Records with missing species identity, temperature or number of workers were excluded.

To assess the factors influencing the success of the Argentine ant at baits, we assembled the species replacement events recorded during the bait surveys. A replacement was defined as the substitution of one species occupying a bait in a given hour by another species in the subsequent hour. For each replacement, we recorded the species involved, the temperature before (T_i_) and after (T_f_) the replacement, and the number of workers of each species. Worker counts were taken for the replaced species in the hour preceding replacement and for the substituting species in the hour the replacement occurred. Each replacement was expressed from the perspective of a focal species against an opposing species, and the success of the focal species (win or loss of the bait) was recorded as a binary variable. Opposing species were classified into four categories: dominant species (excluding the Argentine ant), subordinate species (excluding *Cataglyphis* spp.), *Cataglyphis* spp. (*C. tartessica* and *C. floricola*) and the Argentine ant.

To account for the effect of temperature on the outcome of each replacement, we used the thermal competition index (TCI) proposed by Sánchez-García et al.^45^ The TCI considers both the temperature change during the replacement and how far each species is from its own MAT. First, for each species in each replacement we calculated its improvement rate (ImR) such as: ImR = |T_i_ − MAT| − |T_f_ − MAT| and rounded the result to one decimal place. An ImR value is positive when the final temperature is closer to the MAT of the species than the initial temperature, negative when it is farther from it, and zero when the distance remains unchanged. TCI was then calculated as the difference between the improvement rates of the focal and opposing species. Positive values indicate that the temperature change was more beneficial for the focal species than for the opposing species, and negative values indicate the opposite.

The success of the focal species was modelled with generalised linear models (GLMs) with a binomial error distribution. In the first model, the Argentine ant was the focal species, and the predictors were the category of the opposing species, TCI, and the numbers of workers of both the focal and of opposing species. In the second model, *Cataglyphis* spp. were the focal species, and only replacements involving the Argentine ant were included; thus, opposing species category was omitted as a predictor. Replacements with missing values in any of the model variables were excluded from the analyses. Each coefficient was tested with a Wald z-test, and each predictor with a likelihood-ratio test, which provides a single test for categorical predictors with more than two levels. The maximum-likelihood fit of the second model did not converge because of complete separation. Therefore, this model was refitted with Firth’s penalised-likelihood logistic regression,^46,47^ using the function *logistf()* of the R package *logistf* v1.26.1.^48^

To evaluate the joint effect of temperature and numerical advantage, we fitted two additional models for the same sets of replacements (Argentine ant and *Cataglyphis* spp. as focal species). In these models, we included TCI, replaced the numbers of workers of both species with the base-2 logarithm of their ratio (log_2_[focal/opposing workers]), and the category of the opposing species was not included. With this log transformation, a value of 0 indicates equal numbers, each unit corresponds to a doubling of the ratio, and numerical advantages and disadvantages of the same magnitude are symmetric. The model evaluating *Cataglyphis* spp. was also affected by separation and was fitted with Firth’s penalised likelihood. In the models fitted with Firth’s penalised likelihood, each predictor was tested with a penalised likelihood-ratio test.

Following Sánchez-García et al.,^45^ we used the latter two models to predict the probability of success of the focal species for all combinations of five values of TCI (−20, −10, 0, 10 and 20) and five worker ratios (50:1, 5:1, 1:1, 1:5 and 1:50). The 95% confidence intervals for these predictions were computed on the logit scale and back-transformed to probabilities. Because some of these combinations are rare or absent in the field, we grouped the replacements into classes of worker ratio around the predicted ratios, using class limits at the geometric means of consecutive ratios. A prediction was considered supported by the data when its TCI lay within the range of TCI observed in its class, and predictions outside that range are shown as a theoretical approach.

### 2.6 The effects of treatment in ant community co-occurrence

Co-occurrence analysis tests whether observed patterns of species occurrence differ from those expected under a null model of non-interactions, both spatially and temporally. Here we evaluated whether species co-occurrence patterns were affected by the management in the treated site. To analyse species co-occurrence, we constructed separate species presence-absence matrices for each sampled site, using the pitfall trap groups from each site. We tested differences between both sites before (2000-2005) and after 2005 (2008-2018), when management of the fence was implemented in the treated site. Furthermore, because the spatial effect of the human structure is localised, groups of pitfalls were classified into two distance categories based on their proximity to the human structure, close (0 and 10m) and far (30 and 50m) (Fig. 1b).

Presence-absence matrices were tested for non-random patterns of species spatial co-occurrence using the *cooc_null_model()* function with the sim9 algorithm and 1,000 simulated matrices (after a burn-in of 1,000 randomisations) using the R package *EcoSimR* v0.1.0.^49^ For each community, the standardised effect size (SES) was calculated as the difference between the observed C-score and the mean of the simulated C-scores, divided by their standard deviation. Positive SES values indicate species segregation (less co-occurrence than expected by chance) and negative values indicate species aggregation. Significance was assessed with the one-tailed probability that the simulated C-scores were equal to or greater (segregation) or equal to or lower (aggregation) than the observed C-score; SES values above 2 or below −2 approximately correspond to p < 0.05.

### 2.7 Effects of management on ant community structure

Using species presence and abundance data from pitfall traps, we constructed a species abundance matrix. At each site, two groups of pitfall traps at equivalent distances from the human structure were defined as a given ant community.

For each ant community we calculated the total abundance for each species by summing across all pitfall groups. To prevent the overestimation of the most abundant species, the maximum abundance per species was limited to 25 individuals at each group of pitfall traps. For each site, we tested whether ant communities differed between time periods (before vs after the treatment, implemented in 2005) and distance categories (close vs far to human structures; Fig. 1b). We tested the main effects and their interaction by performing a permutational multivariate analysis of variance (PERMANOVA) using the *adonis2* function, based on Bray–Curtis dissimilarities with terms added sequentially and 999 permutations, of the R package *vegan* v2.7-5.^50^

We graphically represented the ant community structure across both sites by performing a non-metric multidimensional scaling test (NMDS) using the *metaMDS()* function of the R package *vegan* v2.7-5.^50^ With this analysis we represented differences in the structure of the communities in a multidimensional space. To select an appropriate number of dimensions (K = 2) we examine the model stress value, choosing always stress < 0.2 according to Clarke & Warwick.^51^ Plots were generated to display sample clustering across time periods (before vs after the treatment) and the distance from the human structures (close vs far).

### 2.8 Recovery of native endemic species following treatment

To monitor the recovery of the ant community at the treated site relative to the untreated site following the treatment, we visually represented annual ant species abundances from pitfall traps and calculated, for each site and distance category, the mean yearly percentage of workers of each species category before and after the management. We grouped species into four species categories (as described in preceding section): dominant species (excluding the Argentine ant), subordinate species (excluding *Cataglyphis* spp.), *Cataglyphis* spp. or Argentine ant. All analyses were performed in R v4.5.0.^44^

## 3 Results

### 3.1 The success of the Argentine ant at baits

Ant foraging activity at baits was surveyed in 2000, 2001 and 2003, prior to treatment. This allowed us to evaluate how temperature and interspecific interactions influenced the foraging activity of the Argentine ant and native ant species. Bait temperatures did not differ significantly between distances from the human structures or between sites (distance: Welch’s t_253.5_ = −1.27, p = 0.205; Wilcoxon W = 12,904.5, p = 0.198; site: t_344.4_ = −1.39, p = 0.166; W = 13,922, p = 0.129; Fig. S1).

Daily bait foraging in both the Argentine and *Cataglyphis* ants was temperature-dependent. Specifically, the Argentine ant exhibited lower foraging activity during the central hours of the day and was more frequent near the human structures than farther away. *Cataglyphis* ants displayed the opposite pattern: they were more frequent at baits during the central hours of the day and farther from the human structures (Fig. 2).

**Figure 2.**
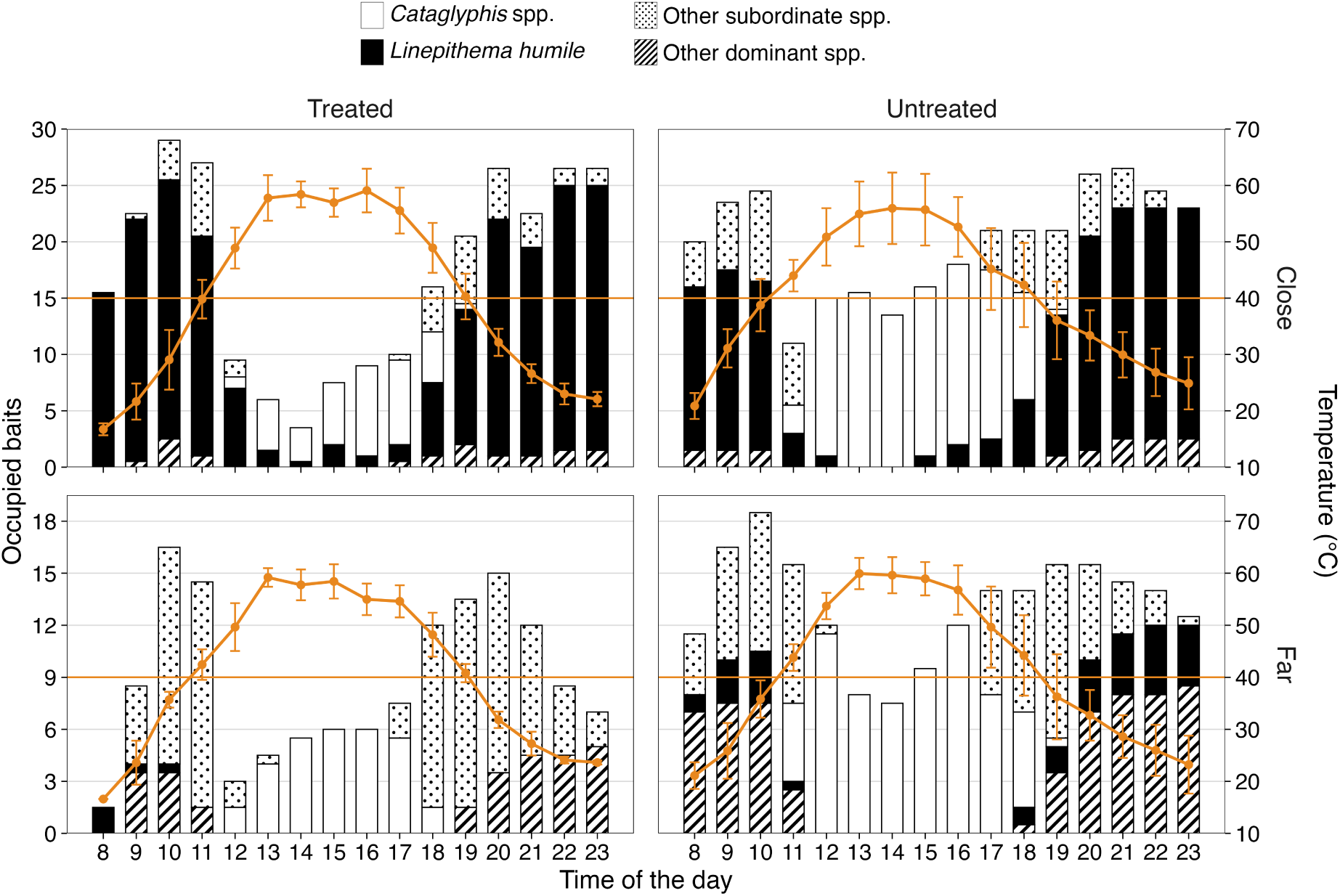
Number of baits occupied by each species category (bars) throughout the day at the two study sites (treated and untreated) and at two distances from the human structures (close and far). Orange points and lines show the mean (± SD) temperature at baits.

The maximum activity temperature (MAT) of the Argentine ant (21.4 °C, n = 756 records) was lower than those of the native species with which it interacted at baits: *Tetramorium forte* Forel, 1904 (30.5 °C, n = 183), *Aphaenogaster senilis* Mayr, 1853 (43.3 °C, n = 446), *C. tartessica* (57.5 °C, n = 403) and *C. floricola* (58.5 °C, n = 62). Across 69 bait replacements with complete data, the Argentine ant won 42 of them: 21 of 35 against *Cataglyphis* spp. (31 replacements with *C. tartessica* and 4 with *C. floricola*), 21 of 33 against the subordinate *A. senilis* and none against the dominant *T. forte*, which displaced it from a bait early in the morning (08:00 h, 16.6–21.0 °C) with 30 workers against 4. Argentine ant success was unaffected by the category of the opposing species (likelihood-ratio test: 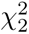 = 1.83, p = 0.400) or by opponent worker numbers (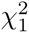 = 0.13, p = 0.716). Instead, success increased significantly with TCI (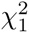 = 54.96, p < 0.001) and with its own number of workers (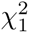 = 5.09, p = 0.024; n = 69; Table S1a). Thus, the Argentine ant was more likely to win when the temperature change favoured it more than its opponent, that is, as temperature approached its low MAT, and when it had more workers.

Of the 35 replacements between *Cataglyphis* spp. and the Argentine ant, *Cataglyphis* spp. won 14, all between 10:00 and 12:00 h and mostly as temperature increased (12 of 14), whereas the Argentine ant won 21, all between 17:00 and 19:00 h and mostly as temperature decreased (20 of 21). *Cataglyphis* spp. success increased significantly with TCI (Firth’s penalised logistic regression; penalised likelihood-ratio test: 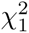 = 25.43, p < 0.001), whereas neither their number of workers (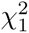 = 0.03, p = 0.854) nor that of the Argentine ant (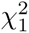 = 2.03, p = 0.154) had a significant effect (whole model: 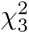 = 34.54, p < 0.001; n = 35; Table S1b). Thus, *Cataglyphis* spp. were more likely to win when the temperature change favoured them, that is, as temperature approached their high MAT, regardless of the number of workers of either species.

When the numbers of workers were replaced by the log_2_ worker ratio, Argentine ant success increased with both TCI (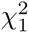 = 58.84, p < 0.001) and its numerical advantage (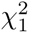 = 4.06, p = 0.044; n = 69; Table S1c), whereas *Cataglyphis* spp. success increased with TCI (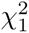 = 29.45, p < 0.001) but not with the worker ratio (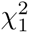 = 2.17, p = 0.141; n = 35; Table S1d). At equal numbers of workers (1:1) and without a thermal advantage for either species (TCI = 0), the predicted probability of success was 0.40 (95% CI = 0.17–0.68) for the Argentine ant and 0.86 (95% CI = 0.35–0.99) for *Cataglyphis* spp. Within the range of conditions observed in the field, both species were very likely to win when TCI was 10 (predicted probability ≥ 0.95 for the Argentine ant and ≥ 0.98 for *Cataglyphis* spp.) and very likely to lose when TCI was −10 (≤ 0.21 and ≤ 0.04, respectively), irrespective of the worker ratio (Fig. 3; Fig. S2 and Fig. S3).

**Figure 3.**
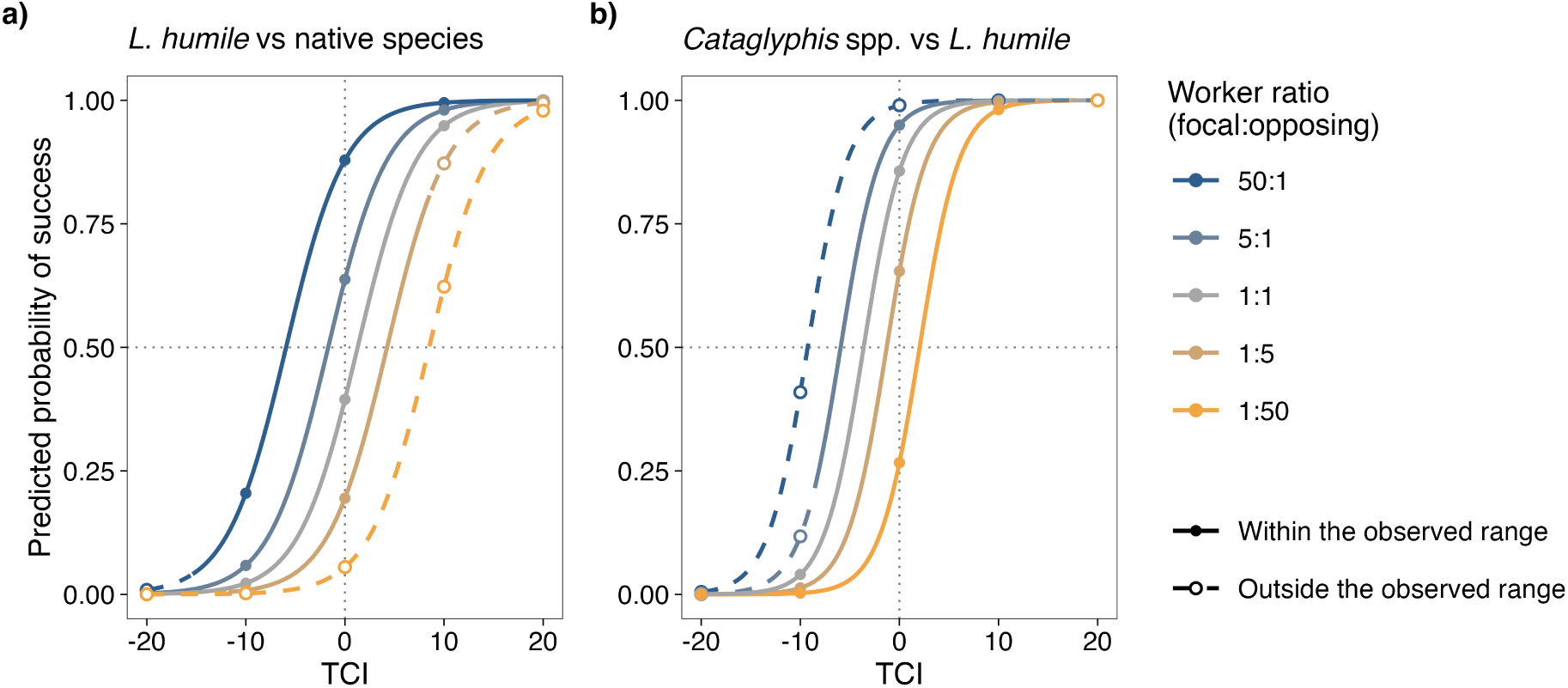
Predicted probability of success in bait replacements as a function of the thermal competition index (TCI) across five worker ratios (focal:opposing species): (a) the Argentine ant against native species and (b) *Cataglyphis* spp. against the Argentine ant. Combinations outside the range observed in the field are represented with dashed lines and open points. Dotted lines mark TCI = 0 and a success probability of 0.5.

### 3.2 Ant community co-occurrence temporal overlap

The co-occurrence analysis tests whether the observed pattern of species occurrence differs from the pattern expected in the absence of interspecific competition, at the spatial and temporal scales. Only one of the eight ant communities exhibited significant species segregation: the community far from the human structure at the treated site after treatment (SES = 3.50, p = 0.001). Although the community close to the human structure at the untreated site before treatment had an SES slightly above 2 (SES = 2.11), its C-score did not differ significantly from random (p = 0.052). The remaining six communities showed a random pattern (SES between−0.79 and 1.50, p ≥ 0.078; Fig. 4).

**Figure 4.**
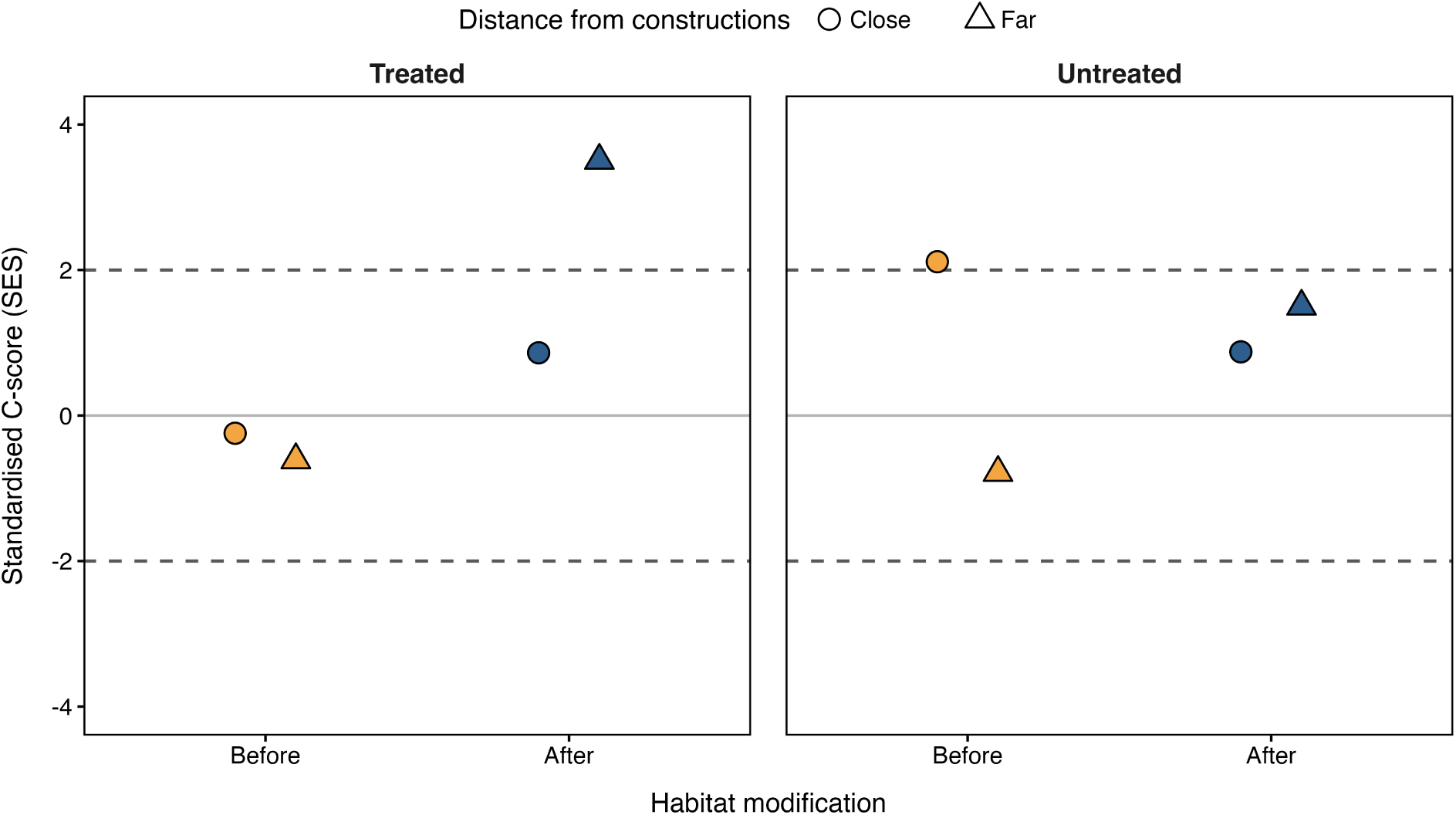
Standardised effect size (SES) of the C-score across study sites (treated and untreated), time periods (before and after treatment), and distances from the human structure (close and far). The dashed lines at SES = ±2 denote approximate limits of statistical significance (p < 0.05).

### 3.3 Ant community structure

PERMANOVA showed that community composition shifted significantly over time only at the treated site (treated: pseudo-F_1,18_ = 6.65, R^2^ = 0.13, p = 0.002; untreated: pseudo-F_1,20_ = 1.13, R^2^ = 0.02, p = 0.316). Distance from the human structure was significant at both sites (treated: pseudo-F_1,18_ = 24.38, R^2^ = 0.48, p = 0.001; untreated: pseudo-F_1,20_ = 26.89, R^2^ = 0.55, p = 0.001), and the interaction between time and distance was not significant at either site (treated: pseudo-F_1,18_ = 1.39, R^2^ = 0.03, p = 0.229; untreated: pseudo-F_1,20_ = 1.18, R^2^ = 0.02, p = 0.277). NMDS ordinations showed clustering by distance to the human structures at both sites, whereas temporal shifts were more evident at the treated site (Fig. 5).

**Figure 5.**
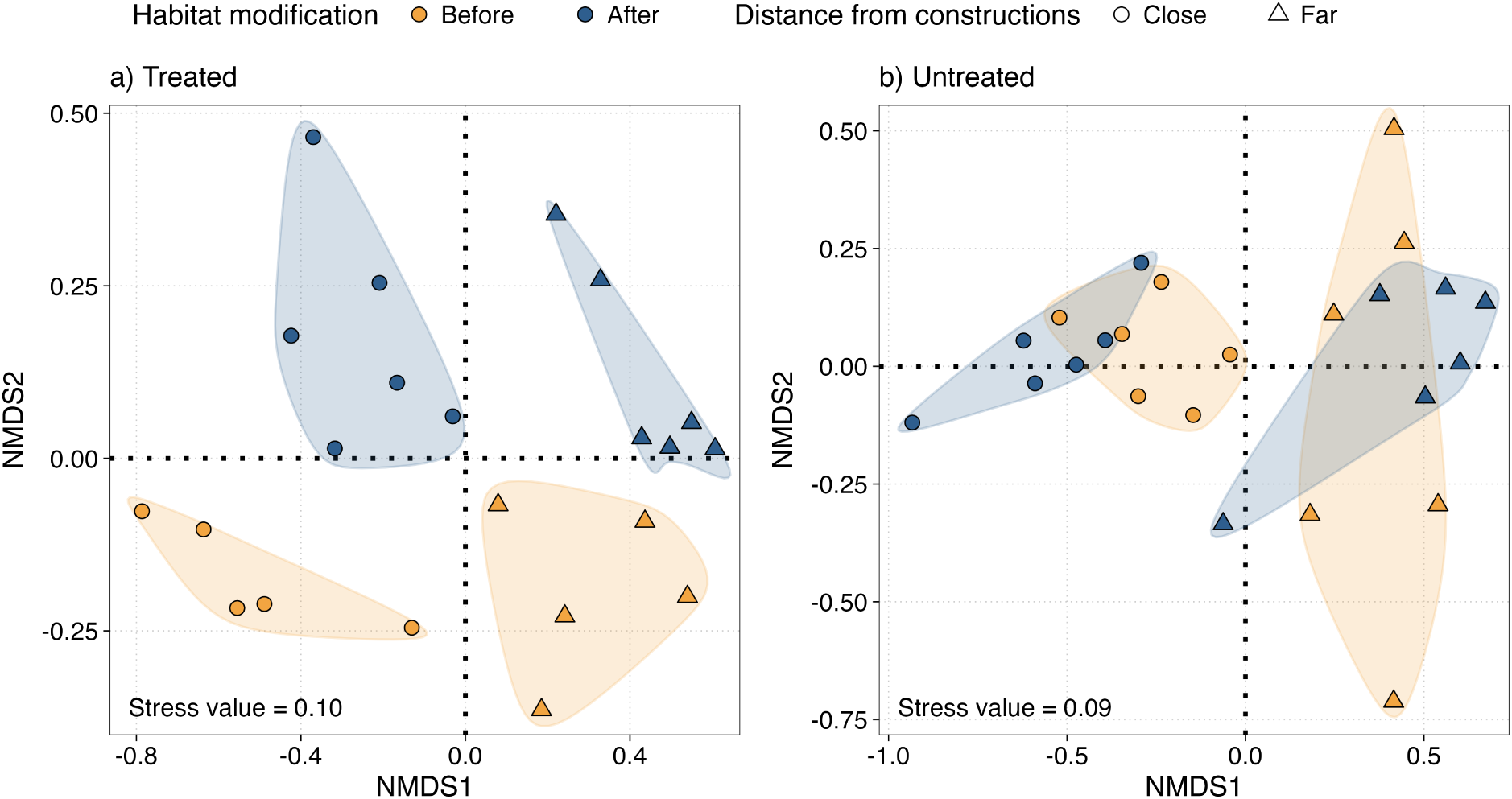
Ordination of ant community similarity (NMDS) across study sites (treated and untreated), time periods (before and after treatment) and distances from the human structure (close and far).

### 3.4 Effectiveness of the treatment on the recovery of native endemic species

Following treatment in 2005, the relative abundance of *Cataglyphis* spp. increased over time at the treated site, but remained low close to the human structure at the untreated site. At the treated site, the mean yearly percentage of *Cataglyphis* spp. among the workers captured increased from 17.8% before to 57.5% after treatment close to the human structure, and from 35.7% to 53.8% far from it, whereas that of the Argentine ant decreased from 64.6% to 22.1% close to the human structure and from 7.1% to 0.2% far from it, where only a few workers were captured after treatment. At the untreated site, close to the human structure, the Argentine ant increased from 77.5% to 90.5% and *Cataglyphis* spp. decreased from 18.3% to 9.0% (Fig. 6).

**Figure 6.**
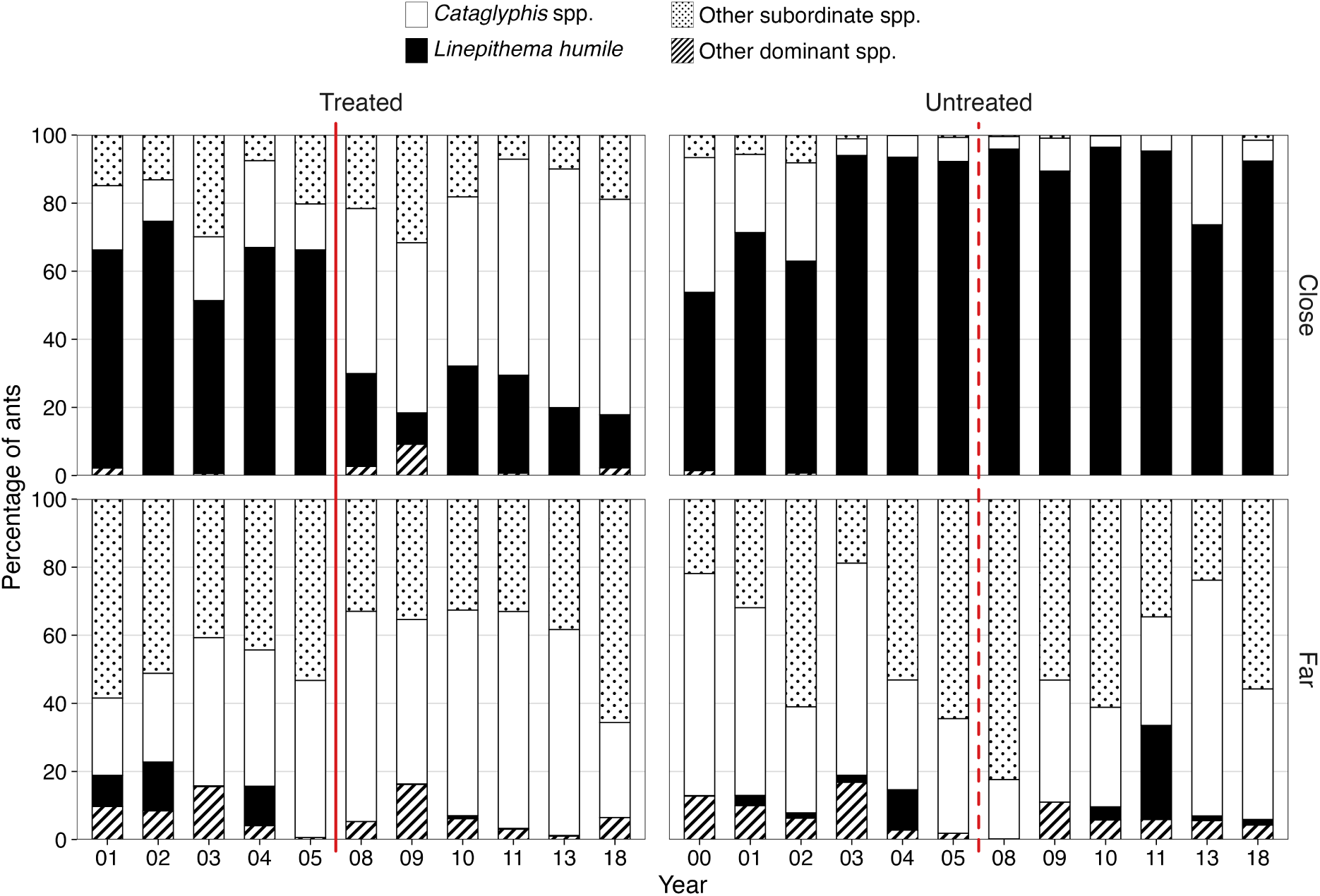
Annual variation in the relative presence of ant species across study sites (treated and untreated) and distances from the human structures (close and far). The red lines separate the years before and after the treatment in 2005 (solid line: treated site; dashed line: untreated site, shown as a reference).

## 4 Discussion

We predicted that management of available optimal habitat could control the Argentine ant invasion. We show that a simple treatment, replacing a concrete fence with a wooden one, reduced the presence and abundance of the Argentine ant at the treated site compared to the untreated site. Ant community structure differed between distances from the human structures, which could be due to differences in microclimatic conditions along this gradient, such as humidity or shade, rather than in temperature at baits, which did not differ significantly between distances. While the Argentine ant succeeded in bait encounters when the temperature change favoured it (i.e., as temperature approached its low maximum activity temperature), the endemic *Cataglyphis* species succeeded when temperature approached their high maximum activity temperature. Significant shifts in ant community structure before and after treatment occurred only at the treated site, where, after treatment, the community far from the human structure became the only one with a segregated pattern of species co-occurrence. Finally, we observed a recovery in the abundance of two endemic ant species, *C. tartessica* and *C. floricola*, at the treated site, especially close to the human structure. However, the treatment was not sufficient to eradicate the invader from this site: 13 years after treatment, the Argentine ant persisted, although at lower abundance.

Native to subtropical South America, the Argentine ant requires suitable microclimates.^32^ It prefers moist habitats with mild temperatures and with dense vegetation, which are not typical of the scrubland of Doñana.^41^ Consequently, the invader colonises this ecosystem primarily near optimal habitat refuges, such as pine forest, cork oak trees or human structures.^40,41,52^ We therefore predicted that replacing a concrete fence – the primary optimal microhabitat at the treated site – with a wooden one, would eradicate the Argentine ant population. However, our results demonstrate that the species can persist at low densities even in the absence of favourable human structures.

Recent studies demonstrate that the Argentine ant can maintain small, non-expanding populations for extended periods, though the underlying mechanisms remain unclear.^39^ Long-term coexistence between Argentine ant and native ant communities has been documented in the Balearic Islands and Corsica, where the Argentine ant fails to achieve complete dominance.^39^ Here, we demonstrate that even after favourable microhabitats are removed, the Argentine ant can persist at low abundance over the long term (13 years), co-occurring with native species far from the human structure that provides optimal abiotic conditions. A similar situation has also been observed in semi-natural habitats in California.^53^

Under the biotic resistance hypothesis, some dominant ant species, such as those of the *Tapinoma nigerrimum* (Nylander, 1856) complex or *Lasius niger* (Linnaeus, 1758), have been proposed to resist Argentine ant invasion.^54–59^ Here we recorded only a single bait replacement (out of 69) involving a dominant ant, *T. forte*, which displaced the invader. No other dominant species interacted with the Argentine ant at baits, probably because the invader is aggressive towards native dominant ants attempting to recolonise the area. In the remaining replacements, the invader was displaced by subordinate species mainly when the temperature change favoured them and when Argentine ant worker numbers were low. This agrees with previous findings in Mediterranean ecosystems, where dominant species win most interactions at moderate temperatures, whereas thermophilic subordinate ants win most interactions at high temperatures, when the dominance hierarchy is disrupted.^60^

Bait encounter outcomes depended mainly on how the temperature change affected each species relative to its own maximum activity temperature. *Cataglyphis* spp. displaced the Argentine ant only in the morning, mostly as temperature increased, whereas the Argentine ant recovered the baits in the late afternoon, mostly as temperature decreased. Numerical advantage mattered only for the Argentine ant, which was more likely to win when it outnumbered its opponent, whereas the success of *Cataglyphis* spp. did not depend on the number of workers of either species. Notably, at a 1:1 worker ratio and without a thermal advantage for either species, *Cataglyphis* spp. were more likely to win than the Argentine ant (predicted probabilities of 0.86 and 0.40, respectively). This suggests that the Argentine ant relies on mass recruitment to displace native species,^16^ whereas the thermophilic *Cataglyphis* can hold baits individually when temperature rises.

In fact, some subordinate species can co-occur with the Argentine ant due to their submissive behaviour, such as *Plagiolepis pygmaea* (Latreille, 1798).^61,62^ In addition, other native ants can co-occur because of different thermal requirements throughout the day, such as the thermophilic and endemic *Cataglyphis* species. Both *C. floricola* and *C. tartessica* have optimal thermal activities around 50 °C^63^ (58.5 and 57.5 °C, respectively, in this study), far above the thermal limit of the Argentine ant (CT_max_: 38–40 °C;^37^ maximum activity temperature of 21.4 °C in this study). This thermal segregation minimises the overlap in their daily activity. However, *Cataglyphis* species nearly disappeared where the Argentine ant was very abundant, at the untreated site close to the human structure, and recovered at the treated site after the treatment reduced the abundance of the invader. In California, Menke et al.^64^ showed that high abundance of the Argentine ant reduced the richness of native species. Here, even the extreme thermal segregation failed to buffer native endemic species against competitive displacement at higher invader abundance. These two *Cataglyphis* species could serve as indicator species, as they are negatively associated with Argentine ant invasion.^65^

Short management actions, such as the treatment presented in this paper, are not enough to control the Argentine ant. The Doñana Biological Reserve has also implemented other measures relying mostly on preventing spread, fostering the clean transport of waste, equipment and experimental material, and of soil or plants.^66^ However, while the Argentine ant spreads using short-range diffusion by budding or long-range dispersal by human transport,^67^ it also benefits from natural vectors such as scavenger raptors.^40^ In this scenario, eradication should be prioritised, together with management actions to prevent spread or control the invasion. Toxic sprays are difficult to implement in a national park because of significant non-target issues,^15^ but other eradication methods (e.g., with granular baits and gels/pastes)^68^ that have had success in other areas should be investigated.^21,69^ Containment and control, together with monitoring and research, should continue to address the long-term effects of invasion and to prevent economic and environmental losses.^19,21^

## Acknowledgements

Many people have helped in the ant surveys along years, such as Aaron Gove, Merav Vonshak, Tahar Slimani, Alain Lenoir, Louise van Oudenhove, Elena Solís, Joan Manuel del Llano, Antonio J. Manzaneda, Fernando Amor, Patro Ortega, Angel Barroso, Ana Carvajal, Irene Villalta and Raphaël Boulay. We thank Jose Manuel Vidal-Cordero for their R programming advice. Fieldwork facilities were provided by ICTS-RBD and we specially thank Rafael Laffitte and the RBD’s staff: Margari, Fabiola and Manolo and all people working at El Palacio facilities since 2000. Ana Carvajal and Isabel Luque helped with sampling separation and identification. This study was funded by the Spanish Ministry of Economy and Competitiveness and FEDER (CGL2015-65807-P) and by Junta de Andalucía (PROYEXCEL_00688) and the Ramón y Cajal Programme (to E.A., RYC-2010-06663).

## Author contributions

**Daniel Sánchez-García:** Data curation, Formal analysis, Investigation, Validation, Visualization, Writing – original draft, Writing – review & editing.

**Xim Cerdá:** Conceptualization, Investigation, Methodology, Writing – review & editing.

**Elena Angulo:** Data curation, Funding acquisition, Investigation, Methodology, Project administration, Resources, Supervision, Writing – original draft, Writing – review & editing.

## Data availability statement

Data and code are archived at Zenodo (https://doi.org/10.5281/zenodo.23105817).

