## Supplementary Figures and Tables for "Temperature and management reduce the abundance of the invasive Argentine ant (*Linepithema humile*)"

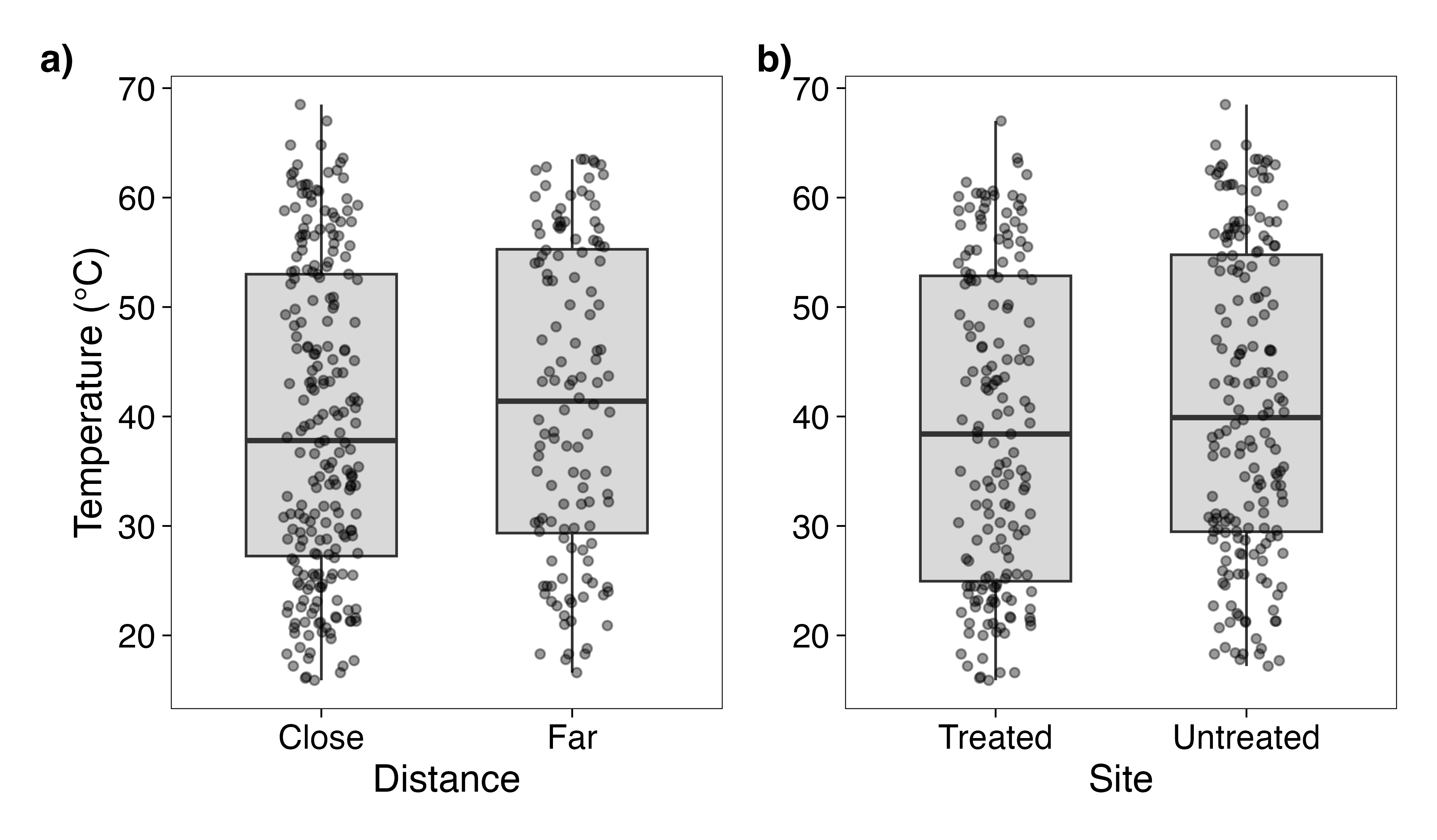

Figure S1. Temperature at baits (a) at two distances from the human constructions (close: plots at 0 and 10 m; far: plots at 30 m) and (b) at the two sites (treated: JL; untreated: MZ), in the bait surveys of 2001 and 2003. Boxes show the median and the interquartile range, whiskers extend to the most extreme values within 1.5 times the interquartile range, and points are the temperature readings (one per plot and hour; n = 351).

Table S1. Logistic regression models of the success of the focal species in bait replacements. Models 1 and 3 (a, c): the Argentine ant (*Linepithema humile*) as focal species against native species (binomial GLMs). Models 2 and 4 (b, d): *Cataglyphis* spp. as focal species against the Argentine ant (Firth’s penalised logistic regression, used because of complete separation). Models 1 and 2 include the numbers of workers of both species, and models 3 and 4 the base-2 logarithm of the ratio between the numbers of workers of the focal and opposing species. TCI: thermal competition index; $\beta$: estimate (logit scale); SE: standard error; n: number of replacements. *p $\leq$ 0.05, **p $\leq$ 0.01, ***p $\leq$ 0.001.

|  | Coefficient (Wald test) | | | | Likelihood-ratio test | | |
| --- | --- | --- | --- | --- | --- | --- | --- |
| Predictor | β | SE | z | p | χ² | df | p |
| **a) Model 1:** Argentine ant vs native species (binomial GLM; n = 69; null deviance = 92.37, df = 68; residual deviance = 23.23, df = 63) | | | | | | | |
| Intercept | −1.21 | 0.90 | −1.34 | 0.181 |  |  |  |
| Opposing species category |  |  |  |  | 1.83 | 2 | 0.400 |
| Dominant species (vs *Cataglyphis* spp.)^a^ | – | – | – | – |  |  |  |
| Subordinate species (vs *Cataglyphis* spp.) | 1.38 | 1.22 | 1.13 | 0.260 |  |  |  |
| TCI | 0.32 | 0.08 | 3.79 | < 0.001*** | 54.96 | 1 | < 0.001*** |
| Workers of the focal species | 0.05 | 0.04 | 1.46 | 0.145 | 5.09 | 1 | 0.024* |
| Workers of the opposing species | −0.04 | 0.11 | −0.39 | 0.696 | 0.13 | 1 | 0.716 |
| **b) Model 2:** *Cataglyphis* spp. vs Argentine ant (Firth’s penalised logistic regression; n = 35; penalised likelihood-ratio test of the model: χ² = 34.54, df = 3, p < 0.001) | | | | | | | |
| Intercept | 1.45 | 1.45 |  |  | 0.91 | 1 | 0.341 |
| TCI | 0.38 | 0.14 |  |  | 25.43 | 1 | < 0.001*** |
| Workers of the focal species | −0.09 | 0.39 |  |  | 0.03 | 1 | 0.854 |
| Workers of the opposing species | −0.03 | 0.02 |  |  | 2.03 | 1 | 0.154 |
| **c) Model 3:** Argentine ant vs native species (binomial GLM; n = 69; null deviance = 92.37, df = 68; residual deviance = 28.03, df = 66) | | | | | | | |
| Intercept | −0.43 | 0.60 | −0.71 | 0.476 |  |  |  |
| TCI | 0.33 | 0.09 | 3.78 | < 0.001*** | 58.84 | 1 | < 0.001*** |
| log_2_ worker ratio (focal/opposing) | 0.43 | 0.24 | 1.76 | 0.079 | 4.06 | 1 | 0.044* |
| **d) Model 4:** *Cataglyphis* spp. vs Argentine ant (Firth’s penalised logistic regression; n = 35; penalised likelihood-ratio test of the model: χ² = 35.64, df = 2, p < 0.001) | | | | | | | |
| Intercept | 1.79 | 1.23 |  |  | 2.18 | 1 | 0.140 |
| TCI | 0.50 | 0.20 |  |  | 29.45 | 1 | < 0.001*** |
| log_2_ worker ratio (focal/opposing) | 0.50 | 0.33 |  |  | 2.17 | 1 | 0.141 |
| ^a^ Not interpretable: this category includes a single replacement (*Tetramorium forte*), so its estimate and standard error are not defined. The other estimates are identical if it is excluded. Models 2 and 4 have no Wald tests, and their tests are penalised likelihood-ratio tests. | | | | | | | |

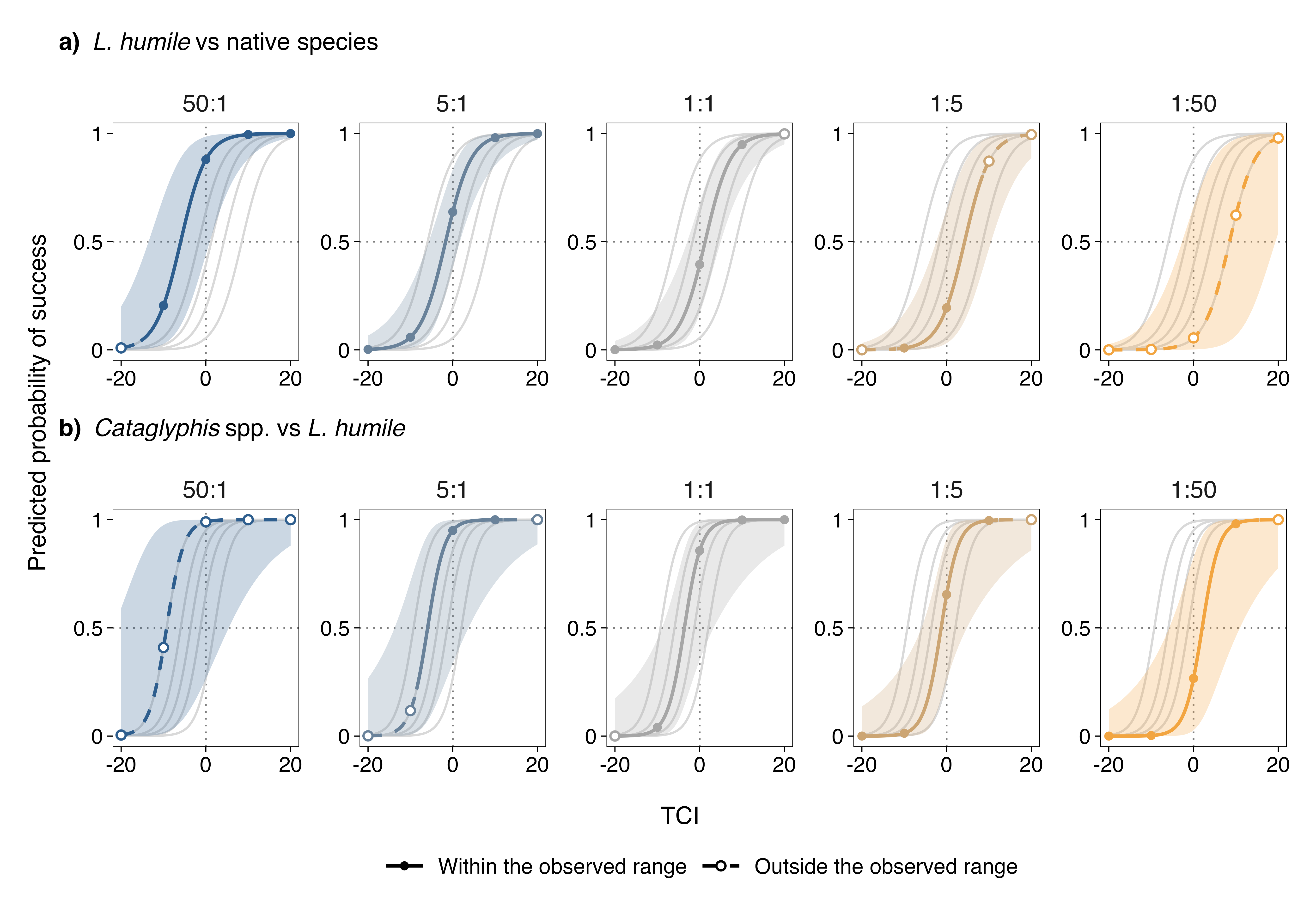

Figure S2. Predicted probability of success in bait replacements along the thermal competition index (TCI), with 95% confidence intervals, for each worker ratio (focal:opposing species): (a) the Argentine ant against native species (model 3, binomial GLM) and (b) *Cataglyphis* spp. against the Argentine ant (model 4, Firth’s penalised logistic regression). Each panel shows the curve of one worker ratio, and the curves of the other ratios in light grey as a reference. Solid lines and filled points show combinations of TCI and worker ratio within the range observed in the field, and dashed lines and open points combinations outside it. Dotted lines mark TCI = 0 and a probability of 0.5.

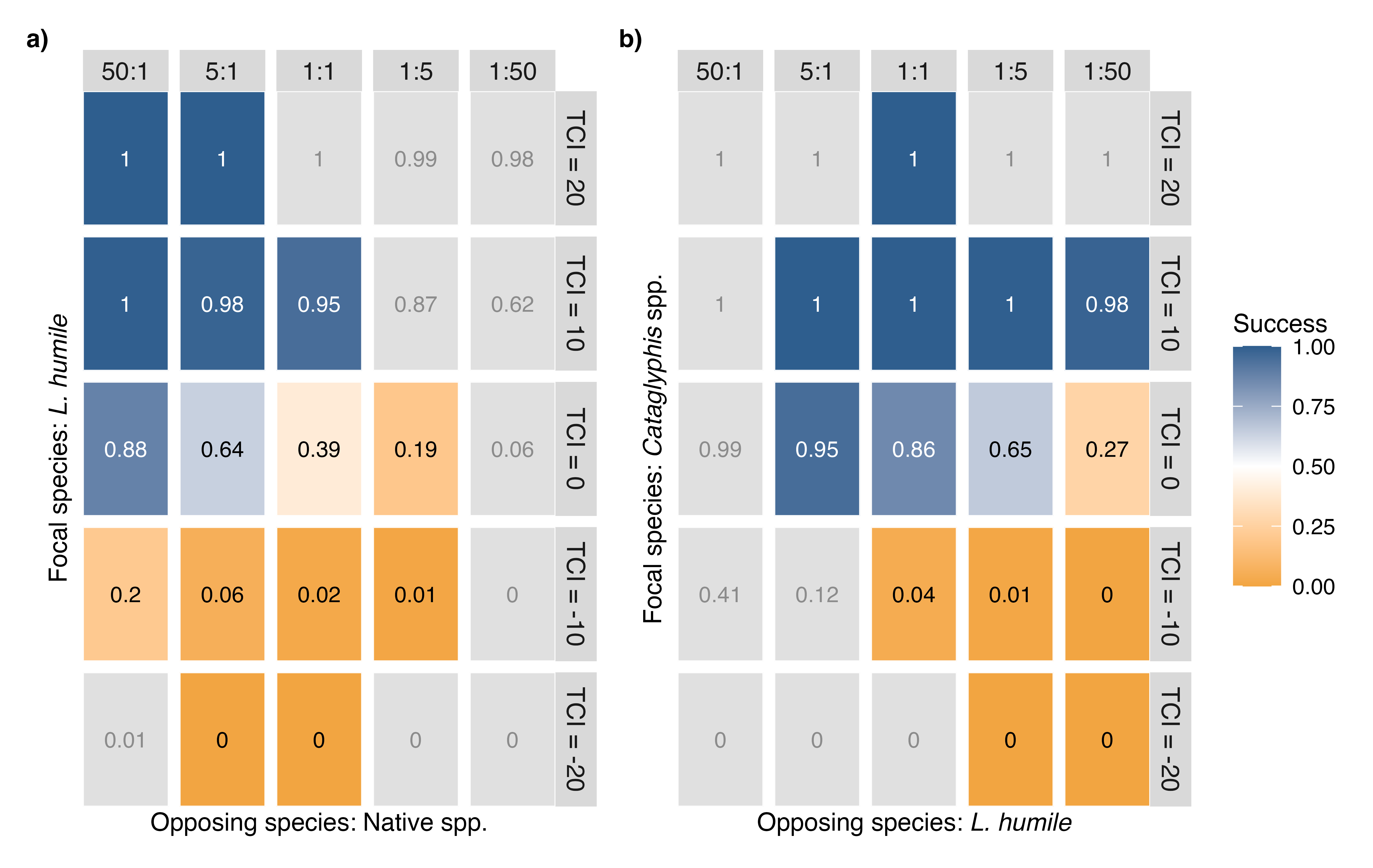

Figure S3. Predicted probability of success (values in the tiles) of (a) the Argentine ant against native species (model 3) and (b) *Cataglyphis* spp. against the Argentine ant (model 4), for five values of the thermal competition index (TCI; rows) and five worker ratios (focal:opposing species; columns). Colours go from orange (0, the focal species always loses) through white (0.5) to blue (1, it always wins). Grey tiles show combinations of TCI and worker ratio not observed in the field, whose predictions are a theoretical approach.
